# Geometric deep learning framework for *de novo* genome assembly

**DOI:** 10.1101/2024.03.11.584353

**Authors:** Lovro Vrček, Xavier Bresson, Thomas Laurent, Martin Schmitz, Kenji Kawaguchi, Mile Šikić

## Abstract

The critical stage of every *de novo* genome assembler is identifying paths in assembly graphs that correspond to the reconstructed genomic sequences. The existing algorithmic methods struggle with this, primarily due to repetitive regions causing complex graph tangles, leading to fragmented assemblies. Here, we introduce GNNome, a framework for path identification based on geometric deep learning that enables training models on assembly graphs without relying on existing assembly strategies. By leveraging symmetries inherent to the problem, GNNome reconstructs assemblies with similar or superior contiguity compared to the state-of-the-art tools across several species, sequenced with PacBio HiFi or Oxford Nanopore. With every new genome assembled telomere-to-telomere, the amount of reliable training data at our disposal increases. Combining the straightforward generation of abundant simulated data for diverse genomic structures with the AI approach makes the proposed framework a plausible cornerstone for future work on reconstructing complex genomes with different ploidy and aneuploidy degrees. To facilitate such developments, we make the framework and the best-performing model publicly available, provided as a tool that can directly be used to assemble new haploid genomes.

## Introduction

*De novo* genome assembly provides essential insights into organism’s biology and evolution by reconstructing its genome from short DNA fragments (or reads), without the access to the original genomic sequence. The assembly process is based on intricate methods on strings and graphs which were first introduced by Esko Ukkonen (Peltola et al. 1984) and later enhanced by Eugene Myers (Myers 1995) and many other scholars. This approach is known as the Overlap-Layout-Consensus (OLC) paradigm, and the initial step is to build an assembly graph where each node represents an individual read, where as pairs of overlapping reads are linked by an edge. The next step, layout, is tasked with placing the reads in the correct order—which is equivalent to finding a path through the graph—and is the core of the genome assembly. An alternative paradigm, based on finding paths in de Bruijn graphs, has been developed shortly after (Idury and Waterman 1995; Pevzner et al. 2001) and these two *de novo* assembly approaches are the most common ones today.

The OLC method does have its challenges. Identifying a path in the graph where each node is visited only once is an NP-complete problem. Furthermore, due to imperfections in the overlap algorithms, certain edges are erroneously missing or falsely present. As a result, much of the research has shifted towards graph simplification, where spurious nodes and edges are eliminated (Myers 2005; Li 2016; Zerbino and Birney 2008; Brankovic et al. 2016; Vaser and Šikić 2021). In many regions, these refined methods yield unique solutions. However, they struggle to handle highly complex, repetitive regions in the graph, and resort to cutting them out. The approaches based on de Bruijn graphs are also often unable to find a unique solution in such regions. Thus, the problem of fragmentation in *de novo* genome assembly persists, unless different types of data and manual curation are used, leading to a slower, more expensive assembly.

Recently, a tremendous achievement has been acccomplished through a combination of both different types of data and manual curation—the complete reconstruction of the human genome (Nurk et al. 2022). This was primarily enabled by the latest advancements in the sequencing technologies, with PacBio HiFi reads and ONT ultra-long reads at the forefront. While the HiFi reads are around 15 kilobases (kb) in length and with error rate of less than 0.5%, the ONT ultra-long reads have error-rates around 5%, but can reach the length of over 100 kb (Li and Durbin 2023). Even though using a combination of these complementary reads achieves state-of-the-art assemblies (Cheng et al. 2023), certain regions of the human genome had to be resolved manually in order to ensure correctness.

This, and similar accomplishments enabled us to develop a novel paradigm for *de novo* assembly, one based on geometric deep learning (Bronstein et al. 2021). Here we present GNNome, a framework which uses an accurate, manually-currated reference genome to produce an arbitrary number of training samples, allowing us to train a model based on graph neural networks (GNNs) (Scarselli et al. 2009) and detect paths in the graph corresponding to genome reconstruction. The contributions of this work are:

- **Contiguity:** While no a-priori knowledge about the algorithmic simplification steps is implemented into the framework, we manage to train a model that yields assemblies with contiguity similar to or higher than the state-of-the-art assemblers across different species on PacBio HiFi data, with some improvements being up to 25% in terms of NG50 and NGA50. We make the best performing model publicly available.
- **Transferability:** The proposed framework is agnostic to the underlying sequencing technology and can adapt to both HiFi and ONT assembly graphs without requiring any modification.
- **Development:** GNNome introduces a shift from traditional C/C++ programming, used in most of the assemblers, to Python/PyTorch-based implementations. This makes the implementation far easier, enabling rapid testing and upgrading with new ideas, fostering faster development cycles. To facilitate the development of new tools, we make the framework’s codebase publicly available.

## Results

### Overview of the GNNome framework

General overview of assembling genomes with GNNome is shown in Fig. 1. It starts with an assembly graph, constructed solely using either PacBio HiFi or ONT reads by an OLC-based genome assembler. Such a graph is passed to a trained model (Supplemental Fig. S1) which assigns a probability to each edge in the graph, reflecting the likelihood that the edge contributes to the optimal assembly. Subsequently, a certain number of edges is sampled as starting points for a search algorithm, which navigates through the probabilities with each walk resulting in a contig (Methods). We demonstrate that a well-trained model can confidently guide a search algorithm to avoid incorrect edges.

**Figure 1.**
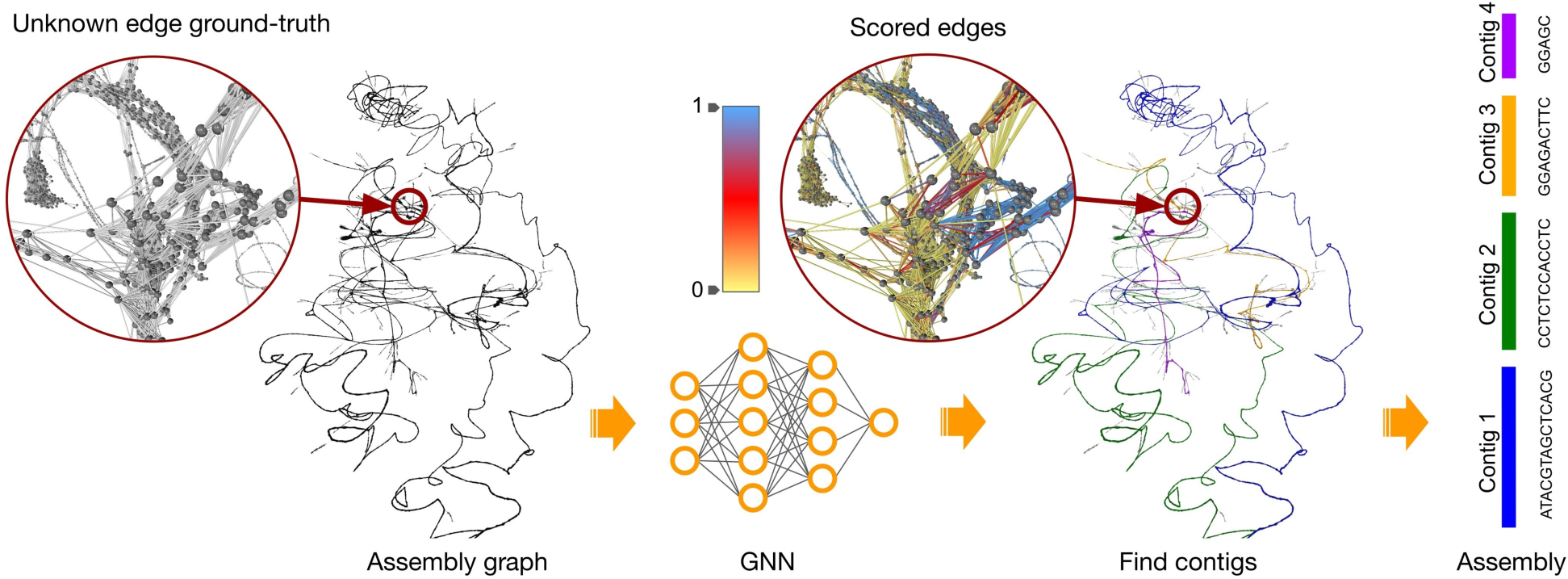
Overview of the assembly workflow with GNNome. An assembly graph is passed to a trained neural network which produces a probability for each edge. The probabilities are visualized in the zoomed-in region in the middle, with the color scheme that increases the visibility of the edges with probability around 0.5. The graph shown here represents entangled chromosomes 21 and 22 of CHM13 (Nurk et al. 2022) generated with hifiasm (Cheng et al. 2021), as well as the four longest contigs found by GNNome. The graph was visualized with Graphia (Freeman et al. 2022).

In conventional assemblers, the assembly graphs are typically simplified with the removal of transitive edges (Myers 2005), trimming of dead-ends (Li 2016), and popping of bubbles (Zerbino and Birney 2008). While these steps are generally reliable and effective, they greatly reduce the number of edges in the graph and thus diminish the amount of information conveyed in the GNN’s message passing. Therefore, we use graphs prior to any simplifications.

The model presented in this study was trained on a dataset constructed from six chromosomes of the human HG002 reference genome (Wang et al. 2022b; Rautiainen et al. 2023) using the PBSIM3 simulator (v3.0.0) (Ono et al. 2022), while the validation dataset, constructed in the same fashion, consisted of five different chromosomes (Methods). The assembly graphs were generated with hifiasm (v0.18.7-r514) (Cheng et al. 2021) as it produces state-of-the-art HiFi-only assemblies, although any OLC-based assembler could be used. Using the saved information about the reads, we implement an algorithm alike breadth-first search (BFS) that determines the label for each edge in the constructed training graphs, allowing for a supervised training of the model (Methods).

Most of the existing algorithms for *de novo* assembly are hand-crafted and their parameters are tuned to several well-known genomes. Usually, it is uncertain whether the good performance transfers to other genomes as well, and fine-tuning the algorithm parameters from scratch for a new set of genomes can be tedious. Here, learning based methods facilitate transferability, as the new genomes can easily be introduced into the training set and the model can be retrained with minimal modifications to the framework.

In this work, we borrow the terminology and the definition of assembly graph from miniasm (Li 2016), where the assembly graph is symmetric since each read is represented with two nodes— one representing the original sequence, and its virtual pair representing the reverse-complement of that sequence (Fig. 2A). Such a definition also ensures that the graph is directed instead of bidirected (Edmonds and Johnson 2003) as some representations are (Myers 2005), which would complicate training of the neural network. To utilize these symmetries during training, we devise a novel GNN layer named SymGatedGCN (Supplemental Fig. S2) and use a loss symmetric with respect to the strand (Fig. 2C). These symmetries were also addressed in several data augmentation steps during the training, such as masking and partitioning of the graph (Fig. 2B) as well as running a search algorithm over the edge probabilities (Fig. 2D). For more details, see Methods.

**Figure 2.**
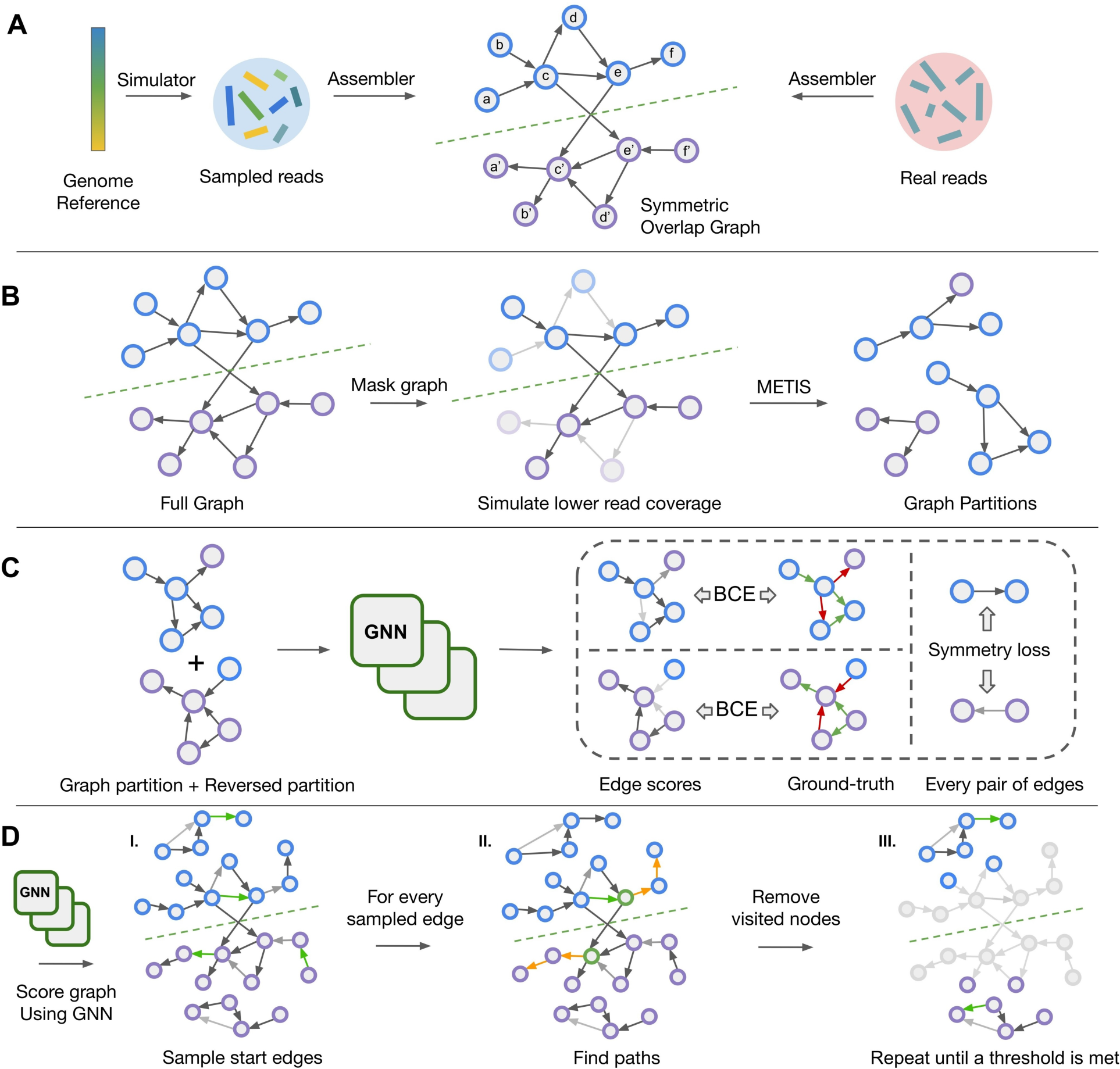
Schematic visualization of the training and decoding pipeline. (*A*) Generating and processing of the synthetic reads for training and real reads for inference. Different colors of the simulated reads indicate that their positions on the reference genome are known, unlike the positions of the real reads. The green dashed line shows the line of mirror symmetry. (*B*) Data augmentation during training. The grayed-out nodes and edges are the ones that are masked. (*C*) Training the model by computing the strand-wise symmetric loss over the edges. The shade of the edges indicates the values of the scores, while red and green color indicates negative/positive edge labels. (*D*) Iterative decoding on a new graph during inference. Green nodes indicate the starting edges for walks, and orange edges represent the steps taken by a decoding algorithm. The grayed-out nodes and edges represent the ones that are previously traversed and are thus discarded in the next iteration of the decoding algorithm.

Contiguity is one of the critical aspects of genome assembly and it is typically assessed using NG50 and NGA50 metrics. Since our methodology is optimized for haploid genome assembly, we evaluate GNNome on the homozygous human genome CHM13 (Nurk et al. 2022) derived from a hydatidiform mole, along with the inbred genomes of *M. musculus* (Hon et al. 2020) and *A. thaliana* (Wang et al. 2022a), all of which are backed by high-quality reference genomes. Additionally, we evaluate on a maternal genome of *G. gallus* (<u>Vertebrate Genome Project</u>), for which the sequencing data was obtained with trio binning.

GNNome is run on the assembly graphs generated by hifiasm (Cheng et al. 2021) from HiFi reads and achieves similar or higher NG50 and NGA50 than the respective assembler. The overview of these results can be seen in Fig. 3. A more detailed analysis, including other measures and the comparison with the state-of-the-art tools as well as transferability to ONT data, is presented in Tables 1 and 2.

**Figure 3.**
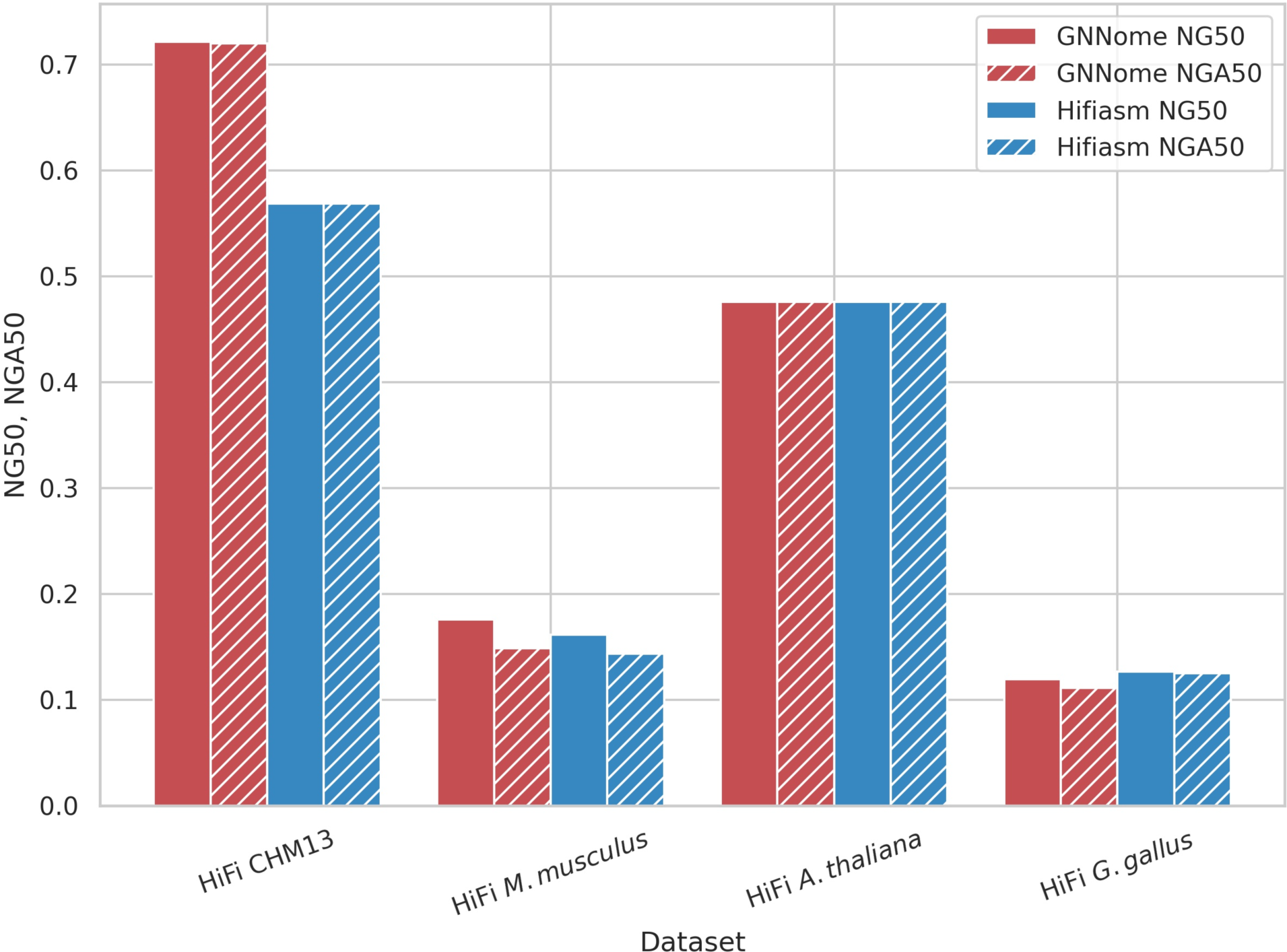
Contiguity on different haploid datasets. Comparison of contiguity of assemblies generated with GNNome and hifiasm (Cheng et al. 2021) for HiFi data. The NG50 of the assembly is defined as the sequence length of the shortest contig that, together with longer contigs covers the 50% of the total genome. The NGA50 is computed similarly like NG50, but instead of considering contigs, we consider blocks correctly aligned to the reference. Both NG50 and NGA50 are presented as proportions of the maximum achievable NG50/NGA50 values for each respective genome.

**Table 1.** Results on HiFi reads. The size is the total length of the assembly. The lengths of the references are 3,054 Mb, 2,728 Mb, 133 Mb, and 1,053 Mb for CHM13 (v1.1), *M. musculus* (GRCm39), *A. thaliana* (Col-XJTU), and *G. gallus* (bGalGal1 maternal) respectively. The LG90 measure is the smallest number of contigs that together cover 90% of the genome. NG50 and NGA50 were computed with minigraph (Li et al. 2020). “Complete” gives the percentage of the reference single-copy genes that are found in the assembly genome, while “duplicated” gives the percentage of reference single-copy genes that are aligned to multiple positions in the assembly. Both “complete” and “duplicated” were computed with compleasm (Huang and Li 2023). QV is per-base consensus accuracy, computed with yak (v0.1-r56) by comparing k-mers in contigs to k-mers found in short reads (Cheng et al. 2021). Short reads were not available for *G. gallus*, so we computed QV with PacBio HiFi reads instead.

| Dataset | Assembler | Size [Mb] | LG90 | NG50 [Mb] | NGA50 [Mb] | Complete [%] | Duplicated [%] | QV |
| --- | --- | --- | --- | --- | --- | --- | --- | --- |
| CHM13 | GNNome | 3,051 | **31** | **111.3** | **111.0** | 99.53 | 0.71 | 54.24 |
|  | Hifiasm | **3,052** | 32 | 87.7 | 87.7 | **99.55** | **0.70** | **55.86** |
|  | HiCanu | 3,297 | 57 | 69.7 | 69.7 | 99.54 | 2.79 | 43.30 |
|  | Verkko | 3,030 | 348 | 9.4 | 9.4 | 99.44 | 0.77 | 51.61 |
| *M. musculus* | GNNome | 2,643 | **140** | **23.0** | **19.3** | **99.62** | 3.30 | 45.40 |
|  | Hifiasm | 2,613 | 150 | 21.1 | 18.7 | **99.62** | **1.93** | 45.67 |
|  | HiCanu | **2,651** | 271 | 11.2 | 10.5 | 99.56 | 2.65 | 43.77 |
|  | Verkko | 2,609 | 204 | 15.9 | 14.6 | 99.60 | 1.95 | **45.72** |
| *A. thaliana* | GNNome | **139** | **13** | **12.4** | **12.4** | 99.89 | 1.09 | **52.08** |
|  | Hifiasm | 151 | **13** | **12.4** | **12.4** | **99.90** | 1.07 | 44.52 |
|  | HiCanu | 152 | 16 | 8.6 | 8.6 | 99.87 | 3.20 | 40.30 |
|  | Verkko | 158 | 18 | 10.3 | 10.3 | 99.87 | **1.04** | 39.75 |
| *G. gallus* | GNNome | 1,114 | 135 | 10.8 | 10.1 | 95.79 | 2.99 | 49.35 |
|  | Hifiasm | **1,087** | **123** | **11.5** | **11.4** | **96.14** | 2.03 | **51.08** |
|  | Verkko | 1,041 | 410 | 3.8 | 3.7 | 95.44 | **1.08** | 49.65 |

**Table 2.** Results on ONT data. The assemblies of GNNome and Raven were additionally polished with Racon (Vaser et al. 2017). Flye and Shasta have built-in polishing modules, so we did not perform additional post-assembly polishing. Shasta did not assemble 90% of the A. thaliana genome, hence the LG90 is undefined.

| Dataset | Assembler | Size [Mb] | LG90 | NG50 [Mb] | NGA50 [Mb] | Complete [%] | Duplicated [%] | QV |
| --- | --- | --- | --- | --- | --- | --- | --- | --- |
| **CHM13** | **GNNome** | **2,984** | **27** | **111.0** | **83.1** | 99.42 | 0.83 | 33.59 |
|  | **Raven** | 2,907 | 48 | 72.6 | 70.3 | **99.51** | 0.70 | 34.04 |
|  | **Flye** | 2,851 | 60 | 70.0 | 69.5 | 99.50 | **0.69** | 33.91 |
|  | **Shasta** | 2,884 | 72 | 59.7 | 56.3 | 99.50 | 0.55 | **37.30** |
| ***A. thaliana*** | **GNNome** | **131** | **9** | **14.6** | **14.5** | **99.83** | 1.07 | 26.54 |
|  | **Raven** | 125 | 11 | 14.5 | **14.5** | 99.81 | 1.07 | **27.28** |
|  | **Flye** | 124 | 15 | 13.7 | 13.7 | 99.79 | 1.07 | 26.03 |
|  | **Shasta** | 115 | - | 12.1 | 12.1 | 99.81 | **0.94** | 25.84 |

### Assembling HiFi datasets

We first evaluate GNNome (v0.4.0) along with hifiasm (v0.18.7-r514) (Cheng et al. 2021), which was used to construct the starting assembly graphs provided to GNNome, on four HiFi samples— CHM13 (Nurk et al. 2022), C57BL/6J strain of *M. musculus* (Hon et al. 2020), Col-0 strain of *A. thaliana* (Wang et al. 2022a), and *G. gallus* maternal genome (bGalGal1 isolate obtained from Vertebrate Genome Project). The coverages of these samples are roughly 32x, 25x, 35x, and 30x, respectively. Additionally, on the same datasets we evaluate HiCanu (v2.2) (Nurk et al. 2020) and Verkko (v1.4.1) (Rautiainen et al. 2023), both popular de novo assemblers. Although hifiasm and Verkko can utilize different types of data in their pipelines, here we focus on HiFi-only assembly.

The results that stand out from Table 1 are the NG50 and NGA50 that GNNome achieves on CHM13—a 25% improvement when compared to hifiasm. It is noteworthy that the model underwent training on graphs of notably smaller sizes—graphs of individual chromosomes further partitioned with METIS (Karypis and Kumar 1998) contained roughly 50 thousand nodes (Methods). Meanwhile, the full graph of the CHM13 genome graphs consists of 2.3 million nodes, showing that the model can generalize well to graphs even 50 times larger and substantially more complex.

Another improvement, though notably less prominent, can be seen in the contiguity of *M. musculus*—GNNome achieved 3% higher NGA50 than hifiasm. In the case of *A. thaliana*, the NGA50 is the same for both methods, and on *G. gallus* GNNome lags behind hifiasm. When evaluating on G. gallus, HiCanu encountered an error after running for 14 days. On the other three datasets, HiCanu achieved lower contiguity by 30% or more. Similarly, Verkko also achieves lower contiguity across the datasets, by more than 15%. It is also worth noting that in case of *A. thaliana*, the length of the GNNome’s assembly is the closest to the length of the reference.

In addition to contiguity, we tested for gene completeness. While the percentage of duplicated genes on CHM13 and *A. thaliana* is similar for GNNome, hifiasm and Verkko, for HiCanu it is considerably higher. The percentage of complete genes is the lowest for Verkko, while for the other three approaches it is similar. On *M. musculus* and *G. gallus*, however, it is GNNome that has more duplicated genes than the other three assemblers. Interestingly, the graph of *M. musculus* seems to be substantially more complex than that of CHM13, even though the *M. musculus* genome is slightly shorter and the coverage of the corresponding read sample was also lower. While the CHM13 graph has 2.4 million nodes and 18.5 million edges, the *M. musculus* graph has 3.7 million nodes and 44.6 million edges. We assume that this higher graph complexity led to lower contiguity and higher percentage of duplicated single-copy genes.

Finally, we compute quality value (QV) to get insight into per-base assembly accuracy. While QV of GNNome’s assembly is slightly lower on CHM13 and *G. gallus* than of hifiasm’s, it is notably higher than QV of all the other assemblers on *A. thaliana*.

We inspect in more detail differences in the CHM13 assembly of GNNome and hifiasm. The most notable difference is in the case of chromosome 11 (Fig. 4). Whereas hifiasm splits a tangle in the centromeric region into three different contigs, GNNome manages to correctly traverse the nodes in the tangle, leading to no breaks and no repetitiveness.

**Figure 4.**
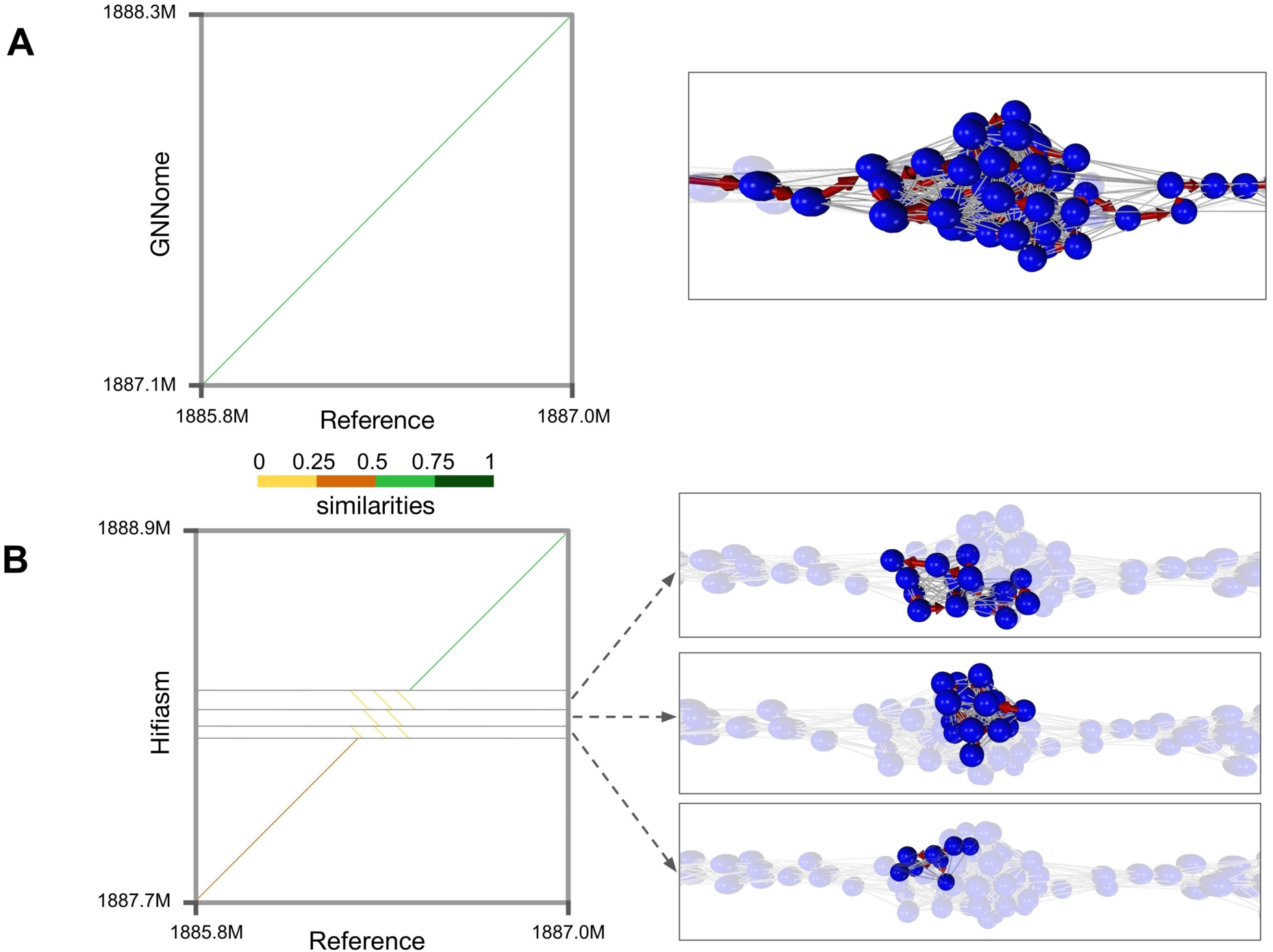
Assembly comparison of chromosome 11 centromeric region. (*A*) GNNome’s assembly of the region. (*B*) Hifiasm’s assembly of the region. Mappings of the assembly to the reference are visualized with dotplots, which were created with D-Genies (Cabanettes and Klopp 2018)—a tool based on minimap2 (Li 2018) alignment. The nodes that comprise each contig are visualized on the right with the help of Graphia (Freeman et al. 2022). The region covered by the tangle is around 264.5 kb long.

### Assembling ONT datasets

We investigate whether the model trained only on HiFi graphs can also generalize to ONT graphs constructed with Raven (v1.8.1) (Vaser and Šikić 2021), without ever seeing any ONT data. Although the idea seems far-fetched at first, both assemblers follow the same sequence alignment principles and construct the graphs in the same fashion, only from the different underlying data. For this purpose, we compare the GNNome’s assemblies of CHM13 and Col-0 strain of *A. thaliana* with Raven’s, starting from the same assembly graphs. The coverage of both samples is roughly 120x. Additionally, we also compare with two other popular assembler for ONT reads, Flye (v2.9.2-b1786) (Kolmogorov et al. 2019) and Shasta (v0.11.1) (Shafin et al. 2020). The results are available in Table 2. Once again, the contiguity of the GNNome, measured by NGA50, on the CHM13 is far higher than in the case of Raven or Flye, and similar to Raven on *A. thaliana*. In addition, the length of the GNN assembly was the closest to the length of the reference genome on both datasets.

It is also worth noting that GNNome assembly of CHM13 achieves substantially higher NG50 than NGA50. This indicates missassembly of certain regions, an issue which was not noticed in HiFi assemblies, likely coming from more accurate edge-predictions on HiFi graphs. These misassemblies are also indicated by the QV and the lower amount of complete single-copy genes found. Such behavior is also not present on *A. thaliana*, where NG50 and NGA50 achieved by GNNome are of similar value, and the number of complete single-copy genes found is higher than for other assemblers.

Although even better results might be achieved by training the model on ONT data from scratch, this approach is not practical from the computational standpoint. The ONT datasets typically have a sequencing depth above 100-fold, considerably higher than the HiFi ones, while also containing sequences of longer lengths. This makes the creating of the read samples, as well as generating the graphs, notably slower. Despite this, the resulting graphs are around two orders of magnitude smaller than the HiFi graphs, primarily due to many reads being discarded as contained. For example, the hifiasm-constructed graph of human chromosome 1 from reads with 60-fold coverage has 311 thousand nodes and 4.1 million edges, while Raven-constructed graph from ONT reads with 120-fold coverage has only 5.6 thousand nodes and 22 thousand edges. Thus, to have the same number of edges in the training set, for each HiFi graph we would require substantially more ONT graphs—making the dataset preparation considerably more computationally expensive.

### Comparing different layout algorithms

It might be assumed that layout algorithms are interchangeable among different genome assemblers, offering little benefit from new implementations. However, when we applied Raven’s layout process to hifiasm-generated graphs, the results contradicted this assumption. According to Supplemental Table S1, the performance of Raven’s algorithm significantly lagged behind hifiasm’s when applied to the same assembly graphs. Specifically, the NG50 and NGA50 metrics for Raven were roughly 50-75% lower than those achieved by hifiasm across all four evaluated genomes, demonstrating the distinct effectiveness of layout algorithms.

This indicates that tailoring assembly algorithms to the particular type of assembly graphs may be present in the existing *de novo* assemblers. Nevertheless, the GNNome model trained only on hifiasm’s graphs achieved high contiguity on Raven’s graph, demonstrating good transferability. This stems from the topological similarity between the two types of graphs, though it also indicates certain differences, leading to imperfections in the assembly of ONT CHM13 dataset.

### SymGatedGCN outperforms other GNN layers

SymGatedGCN layer was inspired by GatedGCN (Bresson and Laurent 2017) due to its performance on several benchmarks (Dwivedi et al. 2020), but additionally includes an edge feature representation and uses a dense attention map for the edge gates (Bresson and Laurent 2019; Joshi et al. 2019). Furthermore, most off-the-shelf GNN layers are devised for undirected graphs. In our earlier efforts (Vrček et al. 2022), we noticed that this is a significant limitation when working with directed graphs. Thus, we devise a bidirectional message-passing procedure with an independent set of trainable parameters for each direction (Methods, Supplemental Fig. S2). Our findings on training on directed graphs were later more formally confirmed (Rossi et al. 2023).

First, we test the effectiveness of SymGatedGCN against a baseline—a model that predicts the same score for every edge, resulting in random walks during the decoding. We demonstrate that random walks vastly underperform when compared to decoding with the model scores (Supplemental Fig. S3).

Next, we compare the performance of SymGatedGCN to other popular GNN layers—GCN (Kipf and Welling 2016), GraphSAGE (Hamilton et al. 2017), GAT (Veličković et al. 2017), and GatedGCN (Bresson and Laurent 2017)—which were run on both directed and undirected graphs, and show consistently superior performance of SymGatedGCN (Supplemental Fig. S4).

Finally, we investigate the influence of node and edge features on contiguity of the assemblies and find that all the features contribute to the performance (Supplemental Fig. S5). There are a few exceptions, such as in the case of HiFi *A. thaliana*, where the model with no node features and the model with no overlap length seem to outperform on average. However, these models greatly underperformed on all the other datasets. This finding highlights the need for expressive node and edge features, suggesting additional feature engineering might further improve the results.

## Discussion

This is the first attempt to solve *de novo* genome assembly using deep learning. Untangling the graph is the central part of the problem, and graph neural networks are perfect for this task— with large amounts of data and complex patterns difficult to recognize, machine learning offers an advantage over engineered algorithms. Moreover, despite being trained only on HiFi graphs constructed with hifiasm, our method can be transferred to other types of data and other assemblers, achieving results comparable or better than other tools. We believe this transferability comes from the similarity in the local structure of both HiFi and ONT graphs— although the graph-construction parameters of hifiasm and Raven are optimized to overlap HiFi and ONT reads respectively, the underlaying logic of both assemblers is the same and is based on sequence alignment algorithms. Interestingly, this does not seem to be the case for the layout algorithms of Raven, a traditional assembler, indicating more general features in our model. Therefore, we believe that our method would bring benefit even to other assemblers that follow the OLC paradigm.

Lately, we have seen the increase in the genomes assembled telomere-to-telomere. Those have mostly been achieved with a combination of different types of data and often manual curation, increasing the time and the cost of the assembly. Improving single-technology assembly leads to not only lower time and cost, but also democratization of assembling new genomes, allowing the laboratories that do not have access to different data types to assemble high-quality draft references.

It is worth noting that all assemblers yield superior results for the human genome compared to other genomes evaluated in this study, likely because of our more comprehensive understanding of human genomics and the greater abundance of such data. This highlights the critical need for intensified efforts toward high-quality genome assemblies of non-human eukaryotic organisms. Similarly, the GNNome model that was trained on a subset of HG002 data achieves the best results on CHM13—both human genomes. As the volume of reliable genome data continues to grow, we anticipate further enhancements in the performance of our deep learning approach, especially on non-human data, and foresee our method serving as a foundation for future approaches designed to leverage this expanding genome dataset.

Drawing inspiration from the pioneering telomere-to-telomere human genome assembly project, which started with a haploid genome (CHM13), our innovative approach also concentrates on haploid assembly. This strategy significantly streamlines the process relative to the assembly of higher ploidy genomes. Moreover, while there exist tools—based on classical algorithms— capable of assembling diploid and even polyploid genomes, they necessitate the use of various sequencing technologies and sequencing of parental genomes to phase the genome reliably. We believe the proposed method can greatly benefit sequence reconstruction when a homozygous or inbred sample is at hand, thereby reducing the need for a large amount of genomic material and cutting sequencing costs, which are often prohibitive. Moreover, the proposed deep learning paradigm, with its capacity to generate abundant simulated datasets and graphs combined with usage of AI methodologies rather than relying on manual inspection and hand-crafted heuristics, is expected to significantly accelerate the development of assembly tools. This advancement will be particularly beneficial for the reconstruction of both diploid and polyploid genomes, as well as in addressing the complexities of aneuploid genomes, where chromosome numbers vary. Such variations are frequently observed in cancerous cells, making this approach highly relevant in cancer research. These more difficult tasks will be the focus of our future work.

We demonstrate the applicability of proposed framework to OLC graphs. However, the same approach can be applied to de Bruijn graphs as well. Moreover, many problems in computational genomics including mapping reads to pangenome graphs belong to combinational optimization problems on graphs which are usually computationally intractable, and the proposed framework can be easily adapted to them.

Finally, even the best layout algorithms are limited by the graphs they are given. The missing edges that were not created during the graph construction cannot be introduced in layout, at least not with the currently available approaches. The increasing popularity of geometric deep learning could open a new frontier here as well—a deep learning-based method for the overlap phase.

## Methods

### Datasets

For training, the data was simulated from HG002 draft reference (version 0.7) (downloaded from <u>HG002 GitHub</u>) and PBSIM3 (Ono et al. 2022). Training set consisted of chromosomes 1, 3, 5, 9, 12, and 18, while the validation set consisted of chromosomes 11, 16, 17, 19, and 20. The acrocentric chromosomes were not considered for either set due to their incompleteness in the draft reference. There was no reason for choosing these exact chromosomes for either training or validation, apart from the fact that we wanted longer and more complex chromosomes in the training set. Moreover, we do not use all the chromosomes from the reference in order to better show the generalization capability of our model to full human genome CHM13 – even though not all the chromosomes were seen in either training or validation, our model consistently performs well. The chromosomes chosen for training were sampled 15 times each to produce samples of reads, which resulted in 15 hifiasm-constructed assembly graphs for each chromosome. Similarly, each chromosome in the validation set was sampled 5 times. Each time the sampling of the reference was different, which led to differences in the assembly graphs used for training and validation. This was a convenient way to synthetically increase the size of our dataset. Note that the graph of each chromosome is considered as a separate data point—the network was not trained on a graph of a combination of reads coming from different chromosomes. For parsing FASTA/Q files, we used Biopython (Cock et al. 2009).

### Labels

The algorithm that determines the label for each edge in the training graphs consists of two steps. In the first step, it utilizes the positional information of each read—start, end, and strand with respect to the reference. It first identifies all edges which are not sampled from the same strand, and which do not have a valid suffix-prefix overlap. More formally, for edge *A* → *B* connecting nodes *A* and *B*, the following relations must hold:

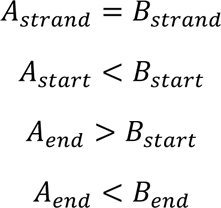

All edges that do not satisfy these criteria are labeled as incorrect and removed from the graph. However, even in such a reduced graph where all the overlaps are valid, not every edge leads to the optimal assembly, the best example being dead ends. Thus, in the second step of the algorithm, a node with the lowest starting position is first identified and BFS is run from it. Of all the visited nodes, the one with the highest end position is picked. Then, we again run BFS, but this time backwards—opposite of the direction of the edges. The edges that are visited by both BFS runs lead to the optimal assembly of that part of the graph and are labeled as positive, while those visited by just one BFS are labeled as negative. In some cases, the assembly graphs can consist of more than one component, so this algorithm is repeated for each of them, until all the edges are labeled. In the implementation of this algorithm, we utilized NetworkX (Hagberg et al. 2008).

The labels we get are highly imbalanced, with around than 99.75% of the edges being labeled as positive. This is expected as hifiasm connects nodes with an edge only if the overlap between the corresponding sequences is of high quality. Thus, most of the false overlaps stem from repetitive regions.

### SymGatedGCN

Let *i* and *s* → *t* be the node and the directed edge whose representations we want to update, respectively. We will use *st* to denote *s* → *t* for simplicity. The node features *x*_*i*_ ∈ ℝ^*d*^_*n*_ and edge features *Z*_*st*_ ∈ ℝ^*d*^_*e*_ are first transformed into their embeddings in the latent space with linear layers. It was shown that, for low-dimensional features, it is beneficial to encode them into a highly dimensional representations in a two-step process (Laurent et al. 2022). In our case, the node features are only in-degree and out-degree of each node, while the edge features are length and the similarity of the overlap, thus *d*_*n*_ = *d*_*e*_ = 2. Similarity is computed as

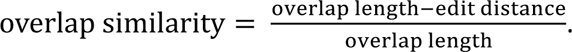

Edit distance computes an approximate string matching between a suffix and the prefix of the overlapping reads (Navarro 2001), and was obtained with Edlib (Šošić and Šikić 2016). The embedding of the features is then performed with:

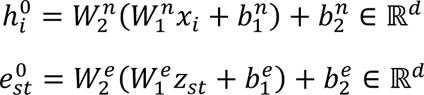

where 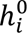 is the initial representation of the node *i*, *e*_*st*_is the initial representation of the edge *st*, and all the *W* and *b* are learnable parameters. The main part of the model consists of a stack of SymGatedGCN layers which update the representations of both the nodes and the edges iteratively. Let the representations of node *i* and edge *st* at layer *l* be 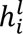 and 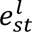. Also, let all the predecessors of node *i* be denoted with *j* and all its successors with *k*. Then, the node and edge representations at layer *l* + 1 will be computed as:

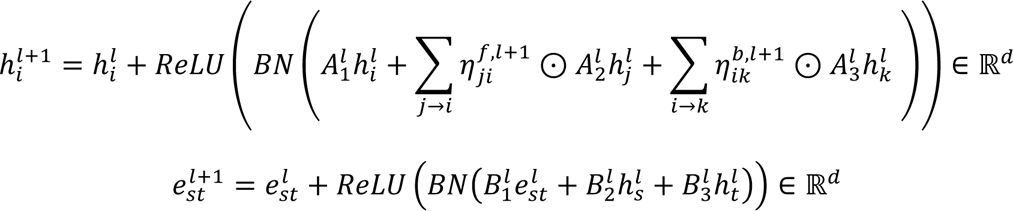

where all *A*, *B* ∈ ℝ^*d*×*d*^ are learnable parameters, *ReLU* stands for rectified linear unit, *BN* for batch normalization, and ⊙ for Hadamard product. The edge gates are defined as:

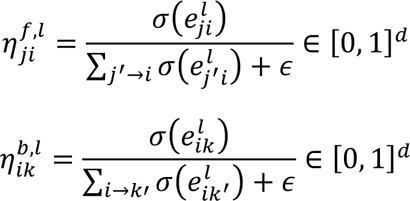

where σ is the sigmoid function, and ε is a small value to avoid division by zero.

Note that we distinguish between the messages 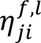 passed along the edges, and in the reverse direction of the edges 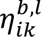. The diagram of this layer, inspired by the diagram of GatedGCN (Dwivedi et al. 2020), can be seen in Supplemental Fig. S2.

Finally, we use a multi-layer perceptron (MLP) on the node and edge representations produced by *L* SymGatedGCN layers to classify the edges. For each directed edge *s* → *t*, a probability *p*_*st*_is computed and trained such that it leads to the optimal assembly. The probability is given by passing the concatenated node representation of nodes *s* and *t*, as well as the edge representation of the directed edge *s* → *t*:

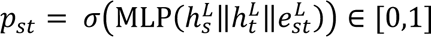

where σ is the sigmoid function, *L* denotes the last SymGatedGCN layer and || is the concatenation operator. The diagram of the whole model can be seen in the Supplemental Fig. S1.

### Data augmentation during training

All the samples of reads were initially generated with a 60-fold coverage, slightly higher than the typical sequencing depth of HiFi reads (30 to 50-fold). This allows us to utilize the reads more efficiently and virtually increase the size of the dataset through masking (Fig. 2B). Every epoch, a subset of nodes of each 60-fold graph is masked to resemble the coverage of between 25 and 60, with the exact number being randomly chosen. The nodes to be masked are also chosen randomly, so each graph can be reused many times as an entirely new data point. Additionally, masking of the nodes is done in a strand-wise manner to retain the symmetry of the assembly graph—either both the node and its virtual pair are masked, or neither of them is.

Since the graphs are simulated from different chromosomes, and then even further reduced while simulating different coverages, their sizes vary greatly. Moreover, training even on the smallest graphs would exceed the graphical processing unit’s (GPU’s) memory. Thus, we partition them into smaller partitions of uniform sizes with METIS (Karypis and Kumar 1998) (Fig. 2B). This ensures that the partitions are roughly uniform in size and that they can be loaded into a GPU memory, which would otherwise be impossible given that even the smallest graphs in the training set consist of approximately 80,000 nodes and 800,000 edges. METIS partitions the graph by cutting the least number of edges needed to achieve a certain number of partitions. Since the edges connecting these partitions are cut out, the model would not make predictions on them during the training, which might hinder the predictions on the similar edges in the inference. Therefore, we add 1-hop neighborhood to each partition to tackle this issue (He et al. 2022).

### Learning objective

The learning objective of the task is binary edge classification, where for each edge the model is predicting whether it leads to the optimal assembly if traversed during the decoding. We utilize the symmetry of the graph and the fact that if an edge *A* → *B* leads to the optimal solution, then necessarily the edge *B*’ → *A*’, connecting the virtual pairs of nodes *A* and *B*, also leads to the optimal solution on the reverse strand (Fig. 2C). More concretely, we devise a loss symmetric with respect to the strand:

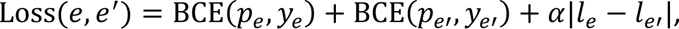

where BCE stands for binary cross entropy, α ≥ 0 is a scalar, *e* and *e*′ are an edge and its virtual pair, whereas *p*, *y*, and *l* denote probability, label, and logit corresponding to the edge *e* or *e*′.

To compute such loss of a certain set of edges, each partition should also include their virtual pairs. This is not guaranteed by METIS. Thus, for each partition a virtual partition is also created, with all the edges reversed, to facilitate the computation of the strand-wise symmetric loss. Those two partitions are considered as a mini-batch, and the backpropagation was performed on the loss averaged over all the edges in the partition (Fig. 2C).

### Training setup

The model used in this study had latent dimension *d* = 64, number of GNN layers *L* = 8, and 3 layers in the MLP classifier. Node and edge features were first linearly transformed to 16-dimensional tensors and then to the 64-dimensional latent space. This results in roughly 220,000 parameters—a small model compared to today’s standards. Dropout of 0.20 was used on node representations after every GNN layer. The scaling factor α in the loss was 0.1. The network was trained with Adam (Kingma and Ba 2014) optimizer with initial learning rate of 10^−4^ and decay of 0.95 after 2 consecutive epochs of no improvement in the validation loss. METIS partitions consisted of roughly 50,000 nodes.

The training was done on a single Nvidia A100 GPU with 40 gigabytes (GB) of memory and took around 9 hours and 15 minutes. For training the network, we used Python (Van Rossum and Drake 2009), PyTorch (Paszke et al. 2019), DGL (Wang et al. 2019), and NumPy (Harris et al. 2020). DGL utilized 32 threads on AMD EPYC 7702 64-Core processor.

### Decoding and inference setup

At inference, METIS is not used in order to avoid cutting any edges in the graph. Thus, we cannot load full graphs into a GPU memory, and instead run inference on a central processing unit (CPU). We perform this on AMD EPYC 7742 processor. A full graph is passed to a trained model, which produces a probability for each edge. The next step is to decode these probabilities and obtain paths representing the genome reconstruction. This is done by sampling the starting edges and running a greedy search from them. Once again, due to the nature of the graph, we opt for the symmetric approach—after sampling a certain number of starting edges, the greedy search in the forward direction is run from the target node of each chosen starting edge, whereas the backward search is run from the virtual pair of the starting edges’ source node (Fig. 2D). The nodes that are not traversed, yet they have a predecessor and a successor that are traversed consecutively, are labeled as visited and discarded—this is equivalent to traversing a transitive edge.

We investigate how the number of starting edges in the decoding influences contiguity. As expected, the higher the number of starting edges, the better the contiguity of the genome, as there is a higher chance that a valid path will be found. However, after increasing the number of starting edges from 100 to 250, on most of the datasets the average results improve only slightly or not at all, as seen in Supplemental Fig. S6. Thus, 100 starting edges gives a good tradeoff between the contiguity and the execution speed (Supplemental Fig. S6). The decoding is performed until the found contigs are longer than 70kb. We noticed that if we keep decoding for shorter contigs, this significantly increases execution time and the number of duplications in our assemblies, stemming from the fact that we use non-reduced assembly graphs.

### Additional analyses

For comparing the performance of the model against random walk, the experiments were performed six times with different random seeds (Supplemental Fig. S3). When comparing the performance of SymGatedCN layer with other GNN layers (GCN (Kipf and Welling 2016), GAT (Veličković et al. 2017), GraphSAGE (Hamilton et al. 2017), GatedGCN (Bresson and Laurent 2017)), the same training setup was used for every run. Each model was trained six times with different random seed and the performance was evaluated on the downstream task, with the same inference setup for every run. For the SymGatedGCN we do the analysis only on the directed graphs, due to the fact that it is implemented in a way to individually handle messages passed in both directions, while for other layers we do the analysis on both the directed (one-direction message passing) and undirected graphs (two-direction message passing). These results can be seen in Supplemental Fig. S4.

Similarly, when investigating the influence of node and edge features on the model’s performance, the same training and inference setups were used, and each model was trained six times with different random seed (Supplemental Fig. S5). For the model that had no node and no edge features, several training runs did not converge. Thus, when running inference, the edge-sampling would fail in some cases due to all edges being assigned a same, near-zero probability. The results here are presented only for the runs that completed the inference.

When investigating the role of the starting edges in the decoding, the edge-probabilities for every run were produced with the same model—the one whose results are presented in Tables 1 and 2. Decoding was run 10 times for every number of starting edges, with different random seed. The results are normalized to 100 starting edges, which is the number we used in all the other experiments (Supplemental Fig. S6).

Finally, we report the time and memory needed for executing the assembly pipeline, both for our method and for other assemblers. The comparison with the other assemblers is not entirely fair, due to the fact that we cannot independently evaluate different parts of their assembly pipelines, such as read overlap, graph construction, and layout. Thus, the time and memory consumption of the other assemblers is provided for their entire pipeline, whereas for our tool we measure each stage independently. Nevertheless, we believe that such analysis gives an insight into the complexity of our approach and the approach of the assemblers, showing that our method is not a bottleneck in the pipeline. These results can be found in Supplemental Table S2. The visualizations described in this section were obtained with Pandas (The pandas development team), Matplotlib (Hunter 2007), Seaborn (Waskom 2021), and Jupyter Notebooks (Kluyver et al. 2016).

### Data availability

All data used in this study is publicly available. For training, we used HG002 draft reference v0.7 which can be found at https://github.com/marbl/HG002. HiFi CHM13 dataset, sequenced with the depth of ∼32x, is available under accessions <u>SRR11292120</u>, <u>SRR11292121</u>, <u>SRR11292122</u>, and <u>SRR11292123</u>. *M. musculus* dataset, sequenced with ∼25x coverage, is available under accession <u>SRR11606870</u>. *A. thaliana* dataset, sequenced with ∼157x coverage and downsampled to ∼35x with SeqKit (Shen et al. 2016) v2.5.1, is available at https://ngdc.cncb.ac.cn/gsa/browse/CRA004538/CRX257575. *G. gallus* HiFi dataset, sequenced with ∼77x coverage was downsampled to ∼30x with Seqkit and is available https://www.genomeark.org/genomeark-all/Gallus_gallus.html. Parental Illumina short reads of *G. gallus*, used in trio-binning, are available on the same link.

ONT CHM13 dataset is available at https://github.com/marbl/CHM13. *A. thaliana* ONT dataset, sequenced with 388x and downsampled to ∼120x with SeqKit, is available at https://ngdc.cncb.ac.cn/gsa/browse/CRA004538/CRX257574. For computing QV, Illumina short reads were downloaded from https://www.ncbi.nlm.nih.gov/sra/?term=SRX1082031 for CHM13, https://ngdc.cncb.ac.cn/gsa/browse/CRA004538/CRX257577 for *A. thaliana*, and from https://www.ncbi.nlm.nih.gov/sra/SRX4381453[accn] for *M. musculus* (Sarsani et al. 2019).

The CHM13 reference (v1.1) can be downloaded from https://s3-us-west-2.amazonaws.com/human-pangenomics/T2T/CHM13/assemblies/chm13.draft_v1.1.fasta.gz.

The *M. musculus* reference GRCm39 (strain C57BL/6J) is available at https://www.ncbi.nlm.nih.gov/datasets/genome/GCF_000001635.27/.

The *A. thaliana* reference Col-XJTU (strain Col-0) can be downloaded from https://download.cncb.ac.cn/gwh/Plants/Arabidopsis_thaliana_AT_GWHBDNP00000000.1/GWHBDNP00000000.1.genome.fasta.gz. The maternal *G. gallus* reference (isolate bGalGal1) is available at https://www.ncbi.nlm.nih.gov/datasets/genome/GCF_016699485.2/.

All the assemblies are available at https://www.dropbox.com/scl/fo/hi5pzdnxhe0jdvjptx0vt/h?rlkey=5pitz9fg3gnbrs9xtlgtnzk5j&dl=0. This is a temporary link, the assemblies will be uploaded to a public Zenodo repository prior to publication.

### Code availability

Source code for GNNome framework (dataset construction, model training, and inference), together with the model weights, is available at https://github.com/lbcb-sci/GNNome.

## Acknowledgements

We thank Robert Vaser and Filip Tomas for help regarding Raven and Racon, especially for implementing functionality that helped our research. We also thank Haoyu Cheng who implemented graph-printing functionality into hifiasm. Finally, we thank Raden Indah Kendarsari for comments on the manuscript.

Lovro Vrček has been supported by “Young Researchers” Career Development Program DOK-2018-01-3373, ARAP scholarship awarded by A*STAR, and core funding of Genome Institute of Singapore, A*STAR. Xavier Bresson has been supported by NUS-R-252-000-B97-133. Martin Schmitz has been supported by SINGA scholarship awarded by A*STAR. Mile Šikić has been supported in part by the European Union through the European Regional Development Fund under the grant KK.01.1.1.01.0009 (DATACROSS), by the Croatian Science Foundation under the project Single genome and metagenome assembly (IP-2018-01-5886), and by the core funding of Genome Institute of Singapore, A*STAR.

## Author contributions

M.Š. and X.B. led the research. L.V., X.B., T.L., and M.Š. developed the neural network architecture, training pipeline, the decoding approach, and curated the data. L.V. implemented the framework, where X.B. contributed to the training and decoding implementation, and M.S. implemented the algorithm for determining the labels of edges in the training graphs. L.V., X.B., and M.Š. devised the experiments, while L.V. performed them and analyzed the results. M.S. composed the figures. M.Š. and K.K. devised the symmetric loss used during the training. M.Š. conceived the initial idea. L.V. and M.Š. wrote the paper, while all the other authors provided feedback.

## Supplemental Material

### Software commands

#### Hifiasm

To generate primary assemblies of homozygous samples from HiFi reads, hifiasm (v0.18.7-r514) was run with the following command:

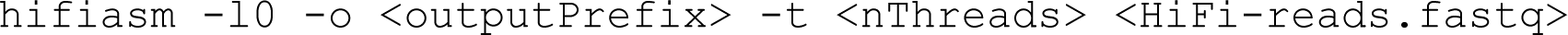

To convert the output GFA files to FASTA files, we ran:

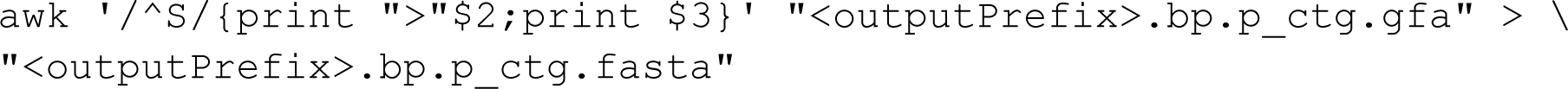

To obtain non-reduced hifiasm graphs which we provided to GNNome, we ran:

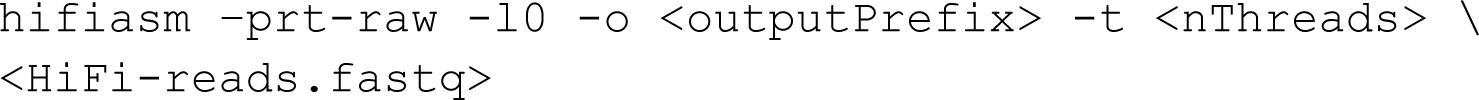

This command was ran to generate graphs of CHM13, *M. musculus*, *A. thaliana* used in the evaluations, but also graphs of individual HG002 chromosomes used during the training.

#### HiCanu

HiCanu (v2.2) assemblies were obtained by running the following command:

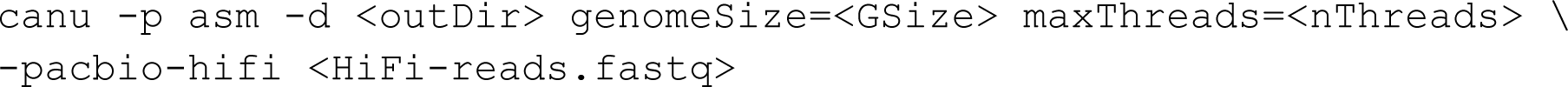

#### Verkko

Verkko (v1.4.1) assemblies were obtained with the following command:

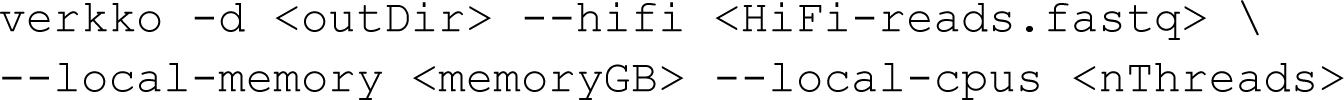

#### Raven

To generate Raven assemblies of ONT data without the polishing step, we ran:

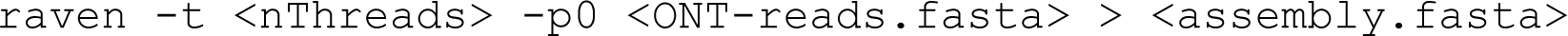

We used Raven v1.8.1 from the print_graphs branch on GitHub (https://github.com/lbcb-sci/raven/tree/print_graphs) which automatically stores the assembly graph in the GFA format to the working directory, under the name graph_1.gfa.

#### Racon

Raven assemblies were additionally polished with Racon. *A. thaliana* assembly was polished with Racon version 1.5.1 for which we first need to generate alignment between the assembly and the reads (PAF file) with minimap2:

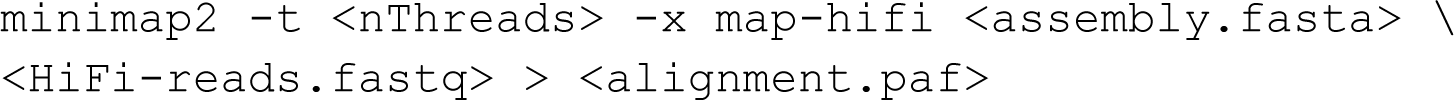

The PAF file, together with the reads and the assembly, can then be passed to Racon to polish the assembly sequence. This was done with the following command:

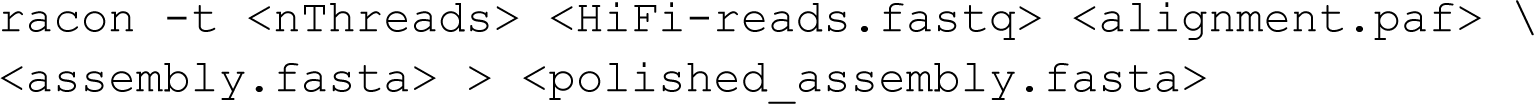

CHM13 was polished with Racon version 3.0.4 which is more lightweight, does not require minimap2 to produce the PAF file, and is integrated into Raven. However, the mappings might be less accurate then with the version 1.5.1 + minimap2. The command used for this was:

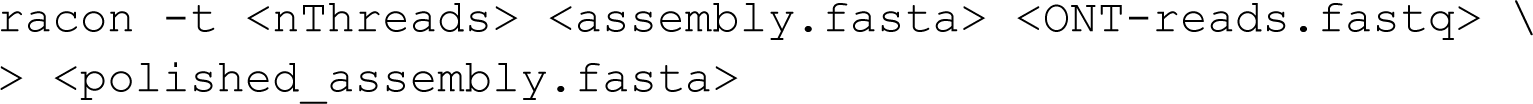

The commands here are for a single round of polishing with Racon. We performed two rounds of polishing, which requires running these commands twice. In the second round

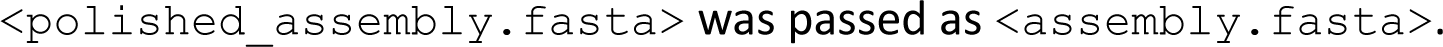

#### Flye

To generate *A. thaliana* assembly with Flye (v2.9.2-b1786), we ran:

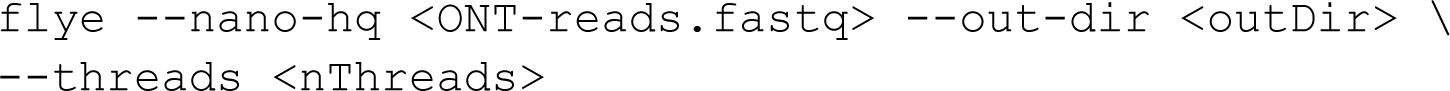

CHM13 Flye assembly was downloaded due to computational constraints.

#### Shasta

To generate ONT assemblies of CHM13 and *A. thaliana* with Shasta (v0.11.1), we ran:

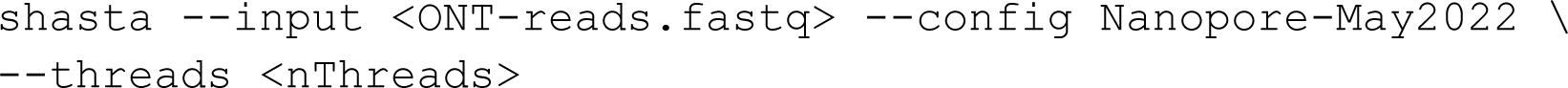

#### GNNome

To obtain GNNome (v0.4.0) assemblies, first step is to parse the GFA files provided by hifiasm and Raven and generate DGL graphs together with auxiliary data (stored in Python dictionaries). For this, we used the following command:

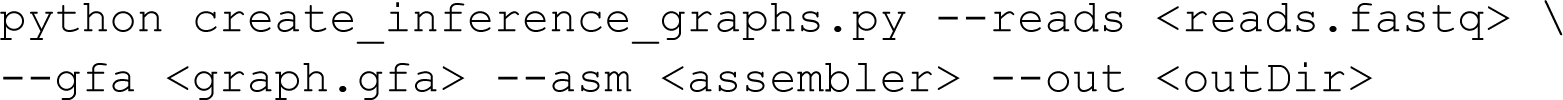

To find the paths comprising the assembly sequence, we ran:

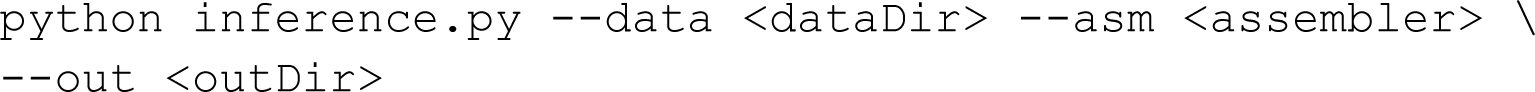

To train the network, we used the following command:

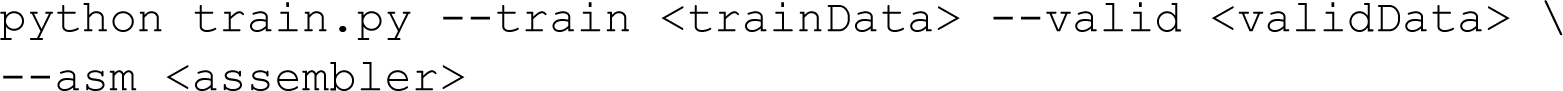

#### Computing NG50 and NGA50

NG50 and NGA50 measures were computed with minigraph (v0.19-r551) and paftools (v2.22-r1101). For running paftools, we used k8 (v0.2.4-r79) JavaScript shell. Minigraph was run with more stringent parameters, as all the reads in this study originate from the same strain as is the reference we are comparing against. This was suggested in the supplementary material of hifiasm. The command used was:

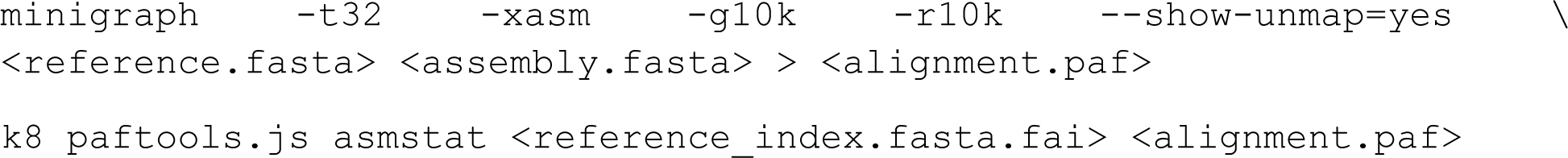

#### Assessing gene completeness

Gene completeness was assessed with compleasm (v0.2.2), using the following command:

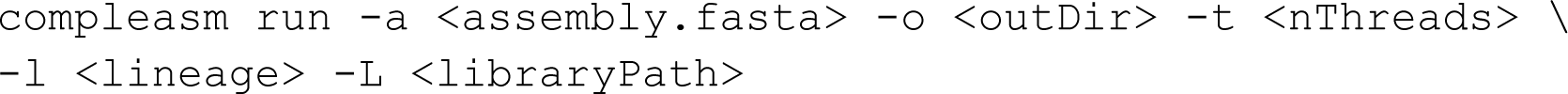

For CHM13, *M. musculus*, and *A. thaliana*,

`<LINEAGE>` was set to primates_odb10, glires_odb10, and brassicales_odb10, respectively.

#### Assembly length and LG90

Assembly size and LG90 were computed in Python, using the Biopython package.

#### Computing QV

Computing QV was done with yak (v0.1-r56). First the kmers were counted, and then the QV was computed with the following commands:

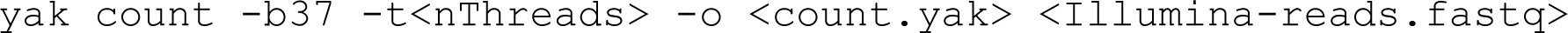

For *G. gallus*, Illumina data of bGalGal1 was not available, so we used PacBio HiFi data

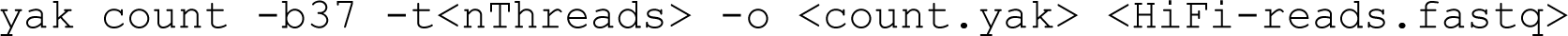

Then, we computed the QV with the following command:

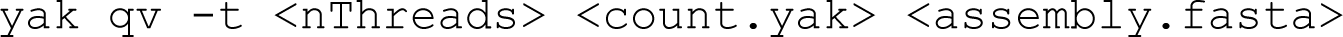

#### PBSIM3

The reads used to construct training and validation graphs were generated with PBSIM3 (v3.0.0). The sample_profile_id was obtained by sampling HG002 HiFi reads. Different chromosomes were sampled individually, with the same sample profile, from the maternal chromosomes of HG002 (v0.7) draft reference. Thus, first the draft reference was split per chromosome. Then, the following command was used to generate the reads:

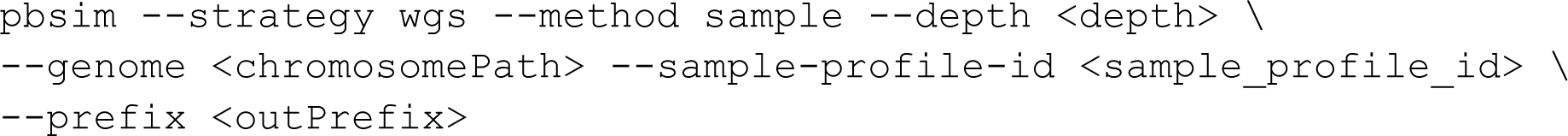

## Supplementary figures

**Supplemental Figure S1:**
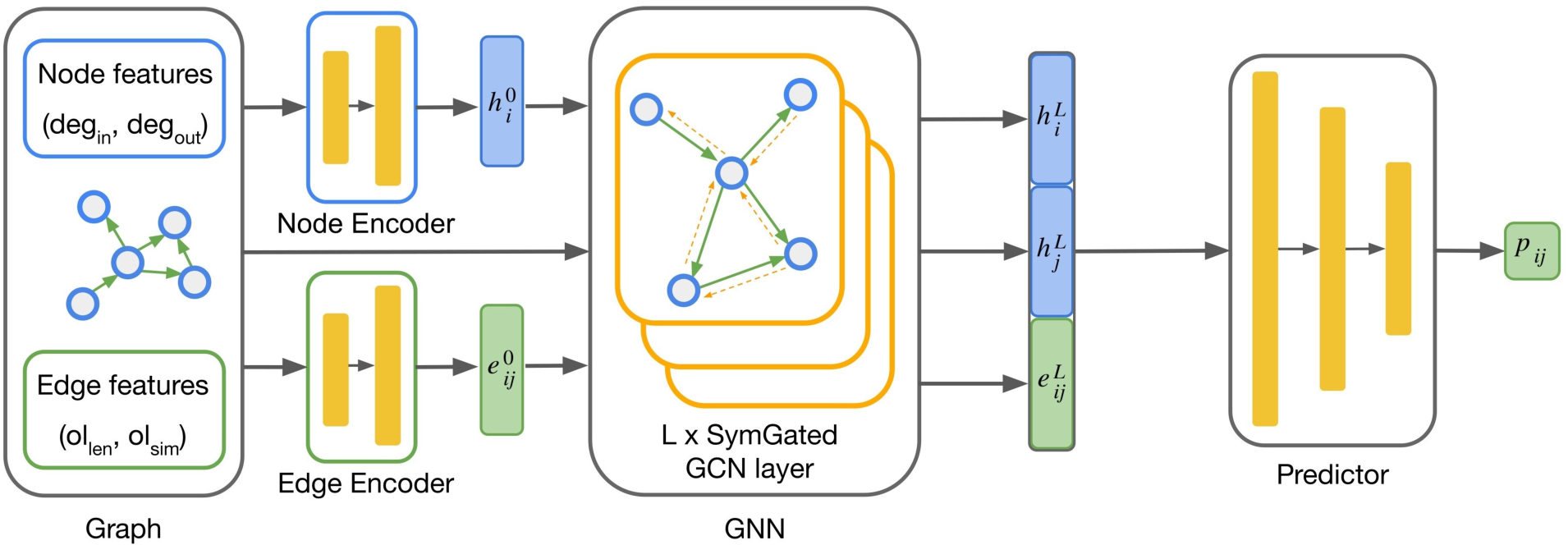
Visualization of the model. Arrows show the flow of information.

**Supplemental Figure S2:**
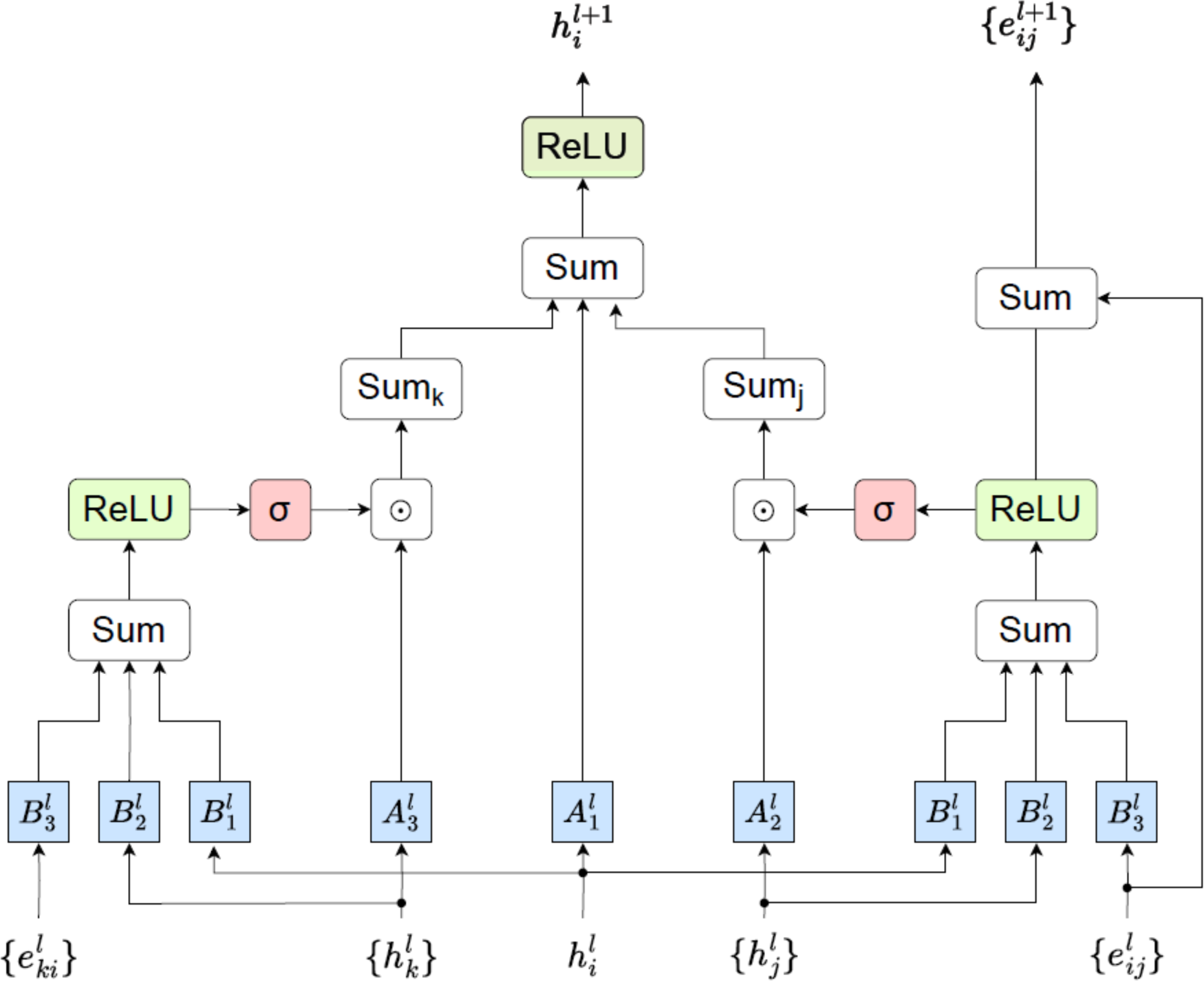
Visualization of SymGatedGCN. The arrows show the flow of information.

**Supplemental Figure S3:**
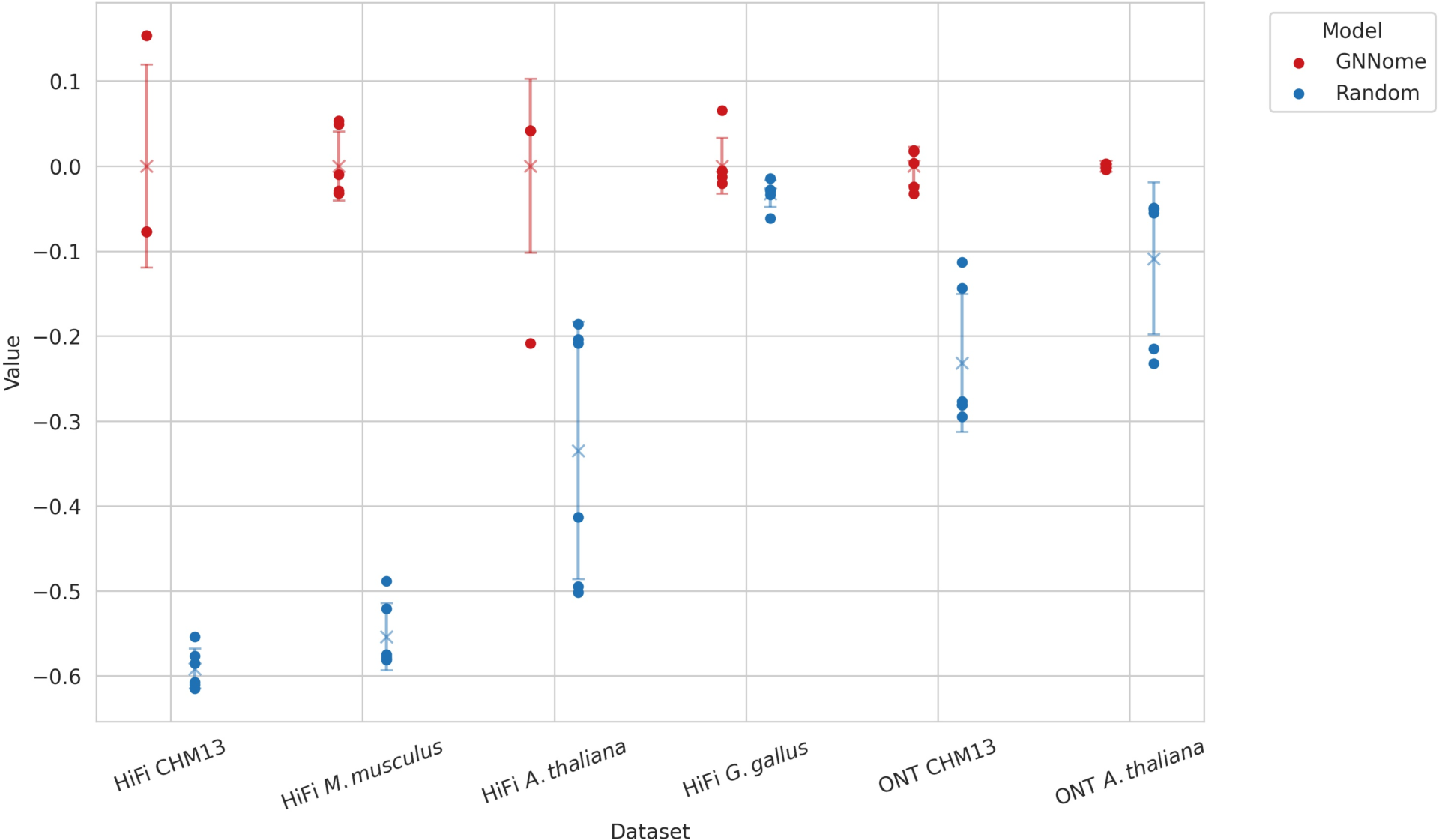
Comparison of GNNome performance against a random walk. The GNNome model was trained six times with six different random seeds, and random walk was also run six times with different random seeds. The result of each run is represented with a dot, while the mean is represented with a cross. The error bars show one standard deviation. The value shown is the mean NGA50, normalized to the mean value of the GNNome.

**Supplemental Figure S4:**
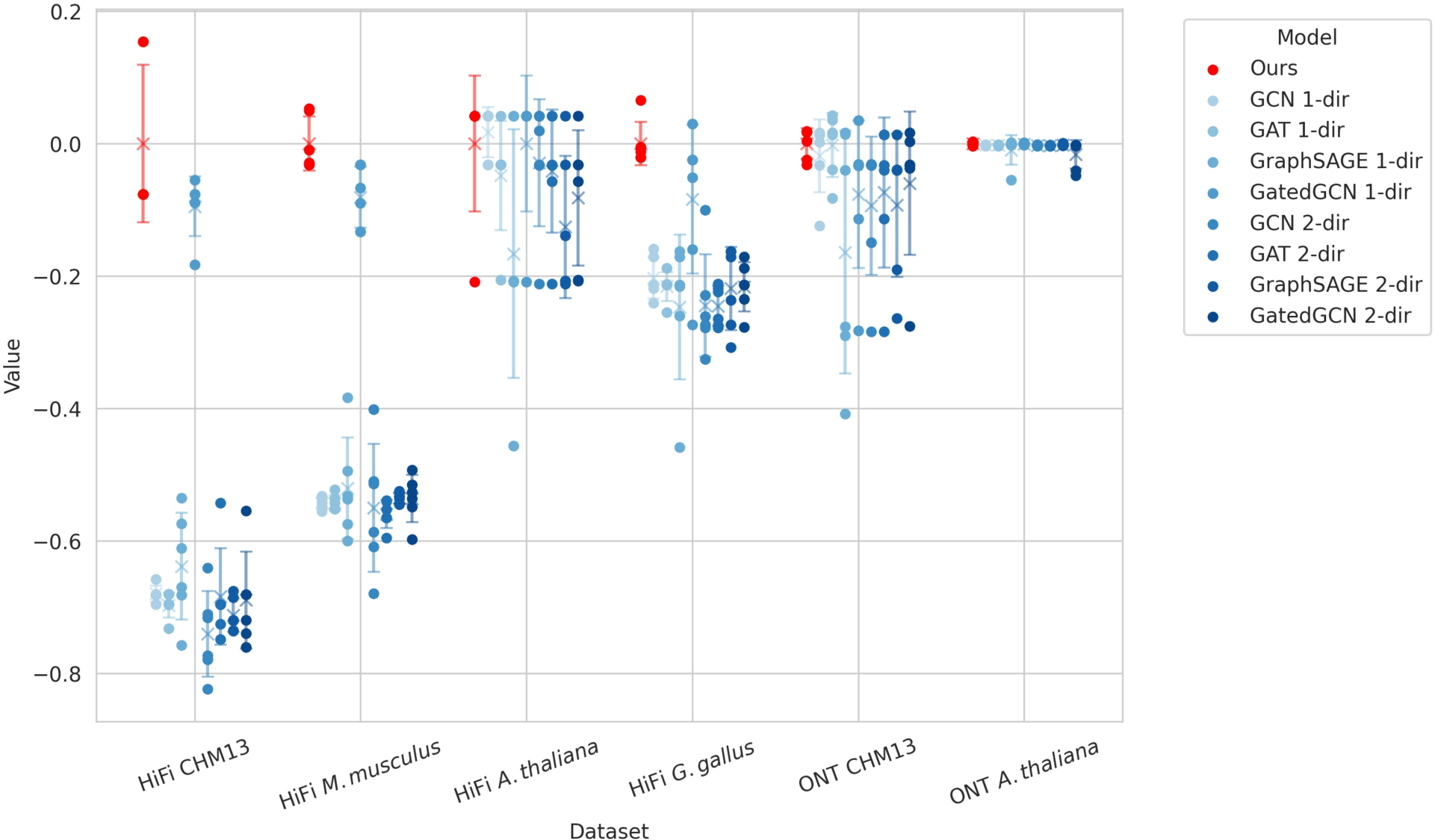
Comparing the performance of different GNN layers. For each layer, the model was trained across six different random seeds. The result of each run is represented with a dot, while the mean is represented with a cross. The error bars show one standard deviation. The value shown is the mean NGA50 of each layer, normalized to the mean value of the best-performing layer (“Ours”). In the legend, “1-dir” stands for one-directional, meaning the message-passing was done on a directed graph, whereas “2-dir” stands for 2-directional, which means that the message passing was performed on an undirected graph.

**Supplemental Figure S5:**
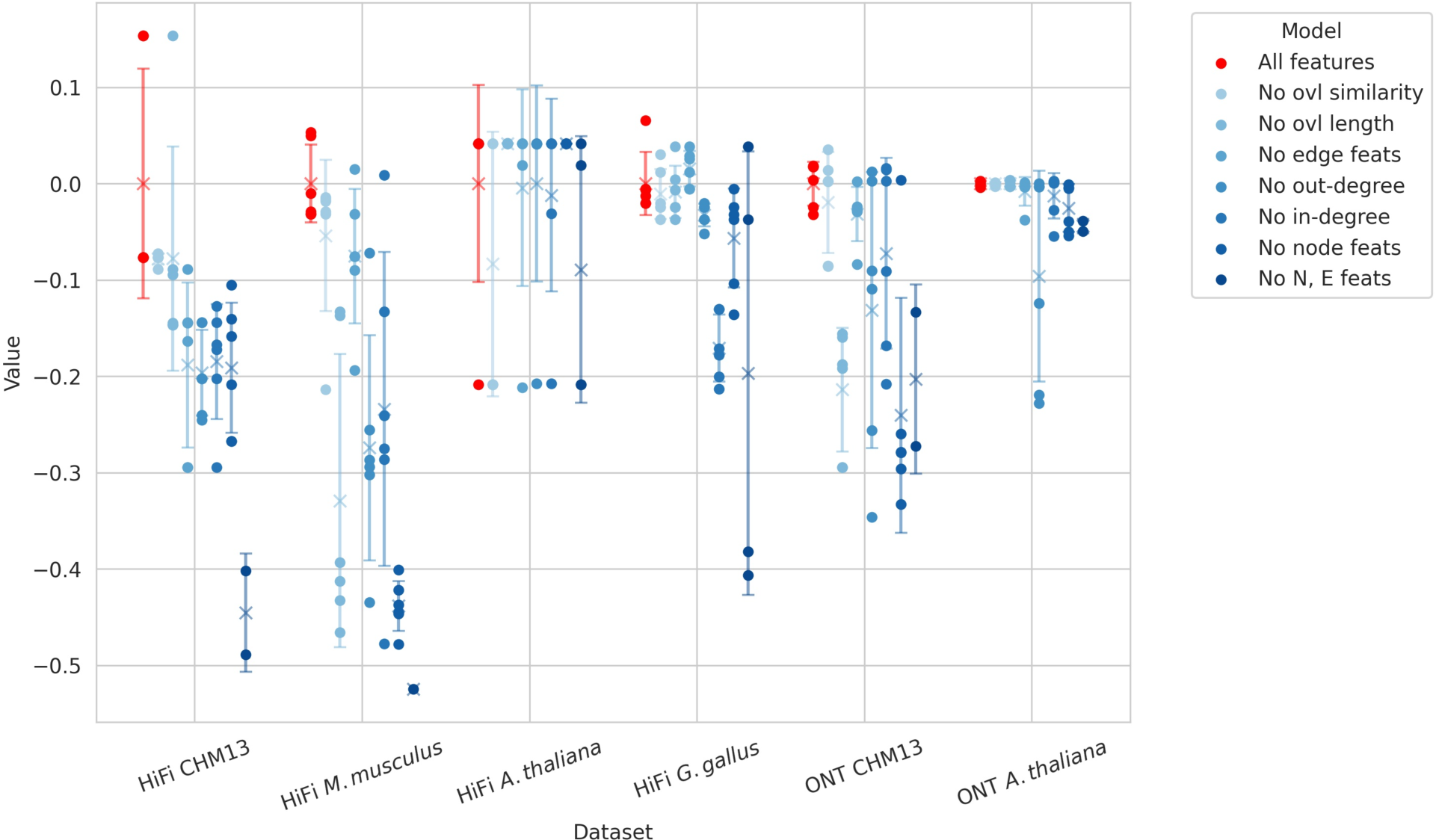
Node and edge feature ablation study. For each set of features, the model was trained across six different random seeds. The result of each run is represented with a dot, while the mean is represented with a cross. The error bars show one standard deviation. The value shown is the mean NGA50 of each set of features, normalized to the mean value of the model with all the features. In the legend, “ovl” stands for “overlap”, “feats” for “features”, “N” for “node”, and “E” for “edge”.

**Supplemental Figure S6:**
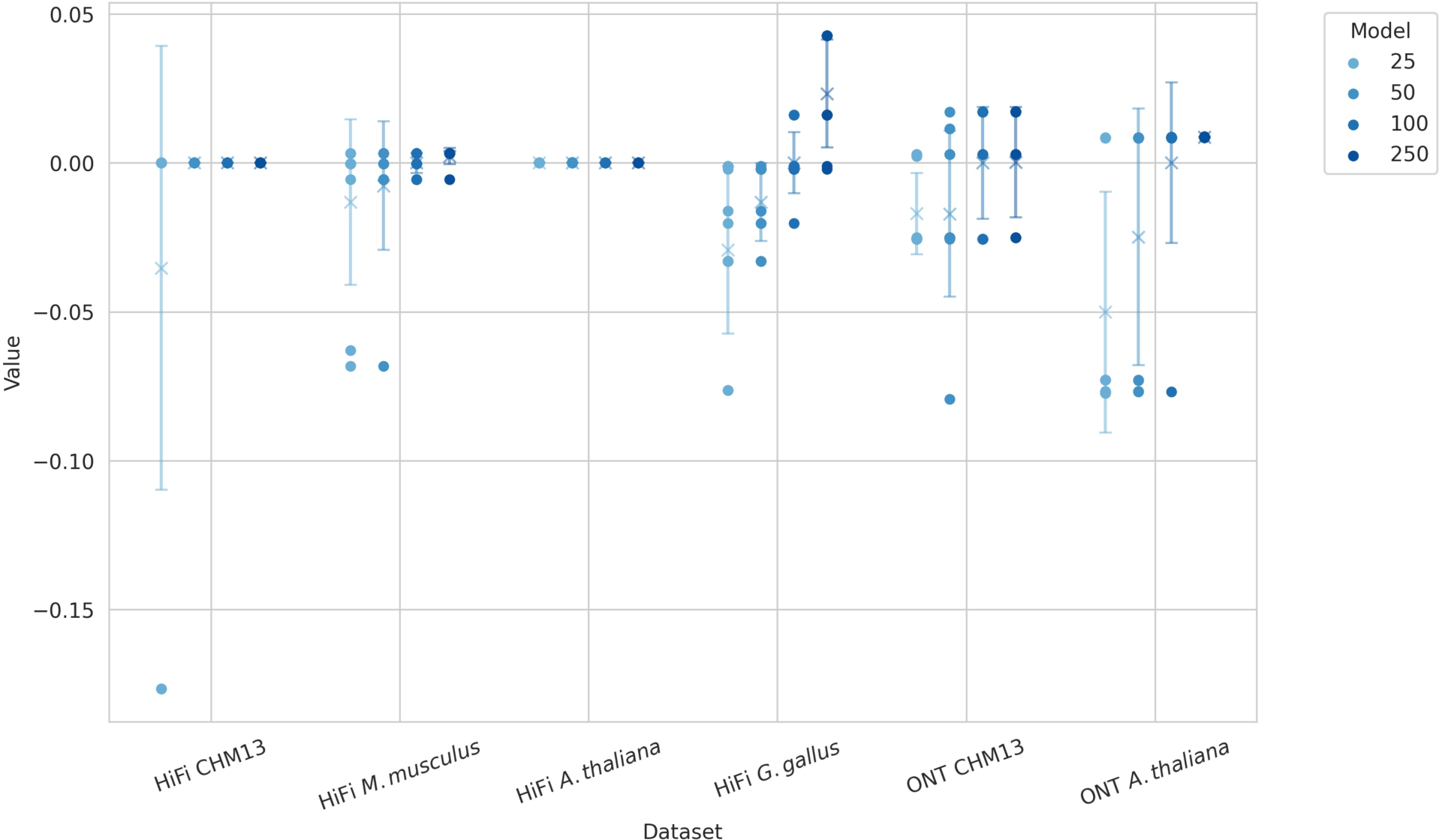
Benchmarking the number of starting edges during the decoding. The model used to produce the edge-probabilities was the best-performing model, whose results are reported in Tables 1 and 2. For each number of starting positions, the decoding was performed with ten different random seeds. The result of each run is represented with a dot, while the mean is represented with a cross. The error bars show one standard deviation. The value shown is the mean NGA50, normalized to the mean value of decoding done with 100 starting edges—the setting used in the rest of the paper (Methods).

## Supplementary tables

**Supplemental Table S1:** Results of Raven’s layout on hifiasm’s assembly graphs.

| Assembler | Dataset | Size [Mb] | LG90 | NG50 [Mb] | NGA50 [Mb] | Complete [%] | Duplicate [%] | QV |
| --- | --- | --- | --- | --- | --- | --- | --- | --- |
| Hifiasm graph + Raven layout | CHM13 | 3,024 | 83 | 45.2 | 45.2 | 99.53 | 0.68 | 49.37 |
|  | *M. musculus* | 2,602 | 253 | 13.1 | 12.9 | 99.59 | 1.89 | 42.69 |
|  | *A. thaliana* | 142 | 14 | 9.4 | 9.4 | 99.82 | 1.02 | 49.49 |
|  | *G. gallus* | 1,076 | 180 | 7.7 | 7.6 | 95.72 | 2.08 | 48.62 |

**Supplemental Table S2:** Comparison of run time and peak memory usage for different assemblers and different parts of the GNNome pipeline. The values for Flye on ONT CHM13 were taken from the paper. All the other values were measured by us. The steps of the GNNome pipeline were performed on an AMD EPYC 7742 processor. The GNN forward pass was performed on 32 threads, while for the rest of the GNNome steps we used a single thread. Other assemblers were run on the same processor with 64 threads. Due to the lack of sudo permissions, Shasta was run with option `--memory-mode anonymous` (default value), which is not optimized for benchmarking (results in higher memory consumprion). The run of HiCanu on *G. gallus* (*) encountered an error after running for the stated amount of CPU time.

| Process | Data type | Dataset | CPU time [h] | Peak RSS [GB] |
| --- | --- | --- | --- | --- |
| **Parse GFA /**<br>**Construct DGL**<br>**graphs** | HiFi | CHM13 | 2.19 | 116.71 |
|  | HiFi | *M. musculus* | 4.69 | 195.13 |
|  | HiFi | *A. thaliana* | 0.10 | 6.68 |
|  | HiFi | *G. gallus* | 0.36 | 25.35 |
|  | ONT | CHM13 | 25.14 | 37.12 |
|  | ONT | *A. thaliana* | 1.94 | 2.52 |
| **GNN forward**<br>**pass** | HiFi | CHM13 | 0.91 | 54.80 |
|  | HiFi | *M. musculus* | 1.54 | 124.11 |
|  | HiFi | *A. thaliana* | 0.06 | 3.61 |
|  | HiFi | *G. gallus* | 0.14 | 8.28 |
|  | ONT | CHM13 | 0.02 | 1.55 |
|  | ONT | *A. thaliana* | 0.01 | 0.43 |
| **Decode /**<br>**Create**<br>**assembly** | HiFi | CHM13 | 1.53 | 60.08 |
|  | HiFi | *M. musculus* | 7.28 | 80.62 |
|  | HiFi | *A. thaliana* | 0.28 | 3.96 |
|  | HiFi | *G. gallus* | 4.41 | 38.3 |
|  | ONT | CHM13 | 0.31 | 28.44 |
|  | ONT | *A. thaliana* | 0.04 | 1.96 |
| **Hifiasm** | HiFi | CHM13 | 318.27 | 116.91 |
|  | HiFi | *M. musculus* | 127.64 | 92.46 |
|  | HiFi | *A. thaliana* | 8.85 | 17.70 |
|  | HiFi | *G. gallus* * | 137.92 | 58.16 |
| **HiCanu** | HiFi | CHM13 | 1206.92 | 96.61 |
|  | HiFi | *M. musculus* | 601.18 | 37.78 |
|  | HiFi | *A. thaliana* | 53.34 | 9.87 |
|  | HiFi | *G. gallus* | 19787.94 | 27.87 |
| **Verkko** | HiFi | CHM13 | 1807.82 | 27.96 |
|  | HiFi | *M. musculus* | 9.23 | 16.56 |
|  | HiFi | *A. thaliana* | 555.72 | 30.33 |
|  | HiFi | *G. gallus* | 94.67 | 21.69 |
| **Raven** | ONT | CHM13 | 2324.32 | 256.64 |
|  | ONT | *A. thaliana* | 4.72 | 21.37 |
| **Flye** | ONT | CHM13 | 4000.00 | 450.00 |
|  | ONT | *A. thaliana* | 80.15 | 91.15 |
| **Shasta** | ONT | CHM13 | 111.00 | 1532.10 |
|  | ONT | *A. thaliana* | 9.21 | 75.62 |

